# Remodelling of cytoskeleton and plasma membrane proteins contributes to drought sensitivity of Arabidopsis *rhd2* mutant

**DOI:** 10.1101/2023.07.11.548511

**Authors:** Tomáš Takáč, Lenka Kuběnová, Olga Šamajová, Petr Dvořák, Jan Haberland, Sebastian T. Bundschuh, Pavel Tomančák, Miroslav Ovečka, Jozef Šamaj

## Abstract

NADPH oxidases are enzymes localised in the plasma membrane and emitting superoxide to the extracellular space. By production of superoxide as one type of reactive oxygen species (ROS), they exert pleiotropic functions in plant development and various stress responses. *Arabidopsis thaliana* RESPIRATORY BURST OXIDASE HOMOLOG PROTEIN C/ROOT HAIR DEFECTIVE 2 (AtRBOHC/RHD2) is an NADPH oxidase with preferential gene expression in roots. Polar localisation and ROS production by this enzyme are essential for root hair elongation. However, the proteome-wide and physiological consequences of *RBOHC/RHD2* mutations are unknown. To find out potential new functions of AtRBOHC/RHD2, we employed a differential proteomic analysis of Arabidopsis *rhd2-1* mutant, carrying a loss-of-function mutation in *RBOHC/RHD2*. Proteomic analyses that were validated with independent biochemical, phenotypical and advanced microscopy methods, showed quantitative deregulation of proteins involved in abiotic and biotic stress response, metabolism, vesicular transport and cell wall modification. Considerable differences in the differential proteomes between roots and above-ground parts were found in the mutant. The altered abundance of aquaporins and homeostasis of transmembrane pumps and transporters most likely determine the higher sensitivity of Arabidopsis *rhd2-1* mutant to drought.

**Highlight:** Proteomics and advanced microscopy reveal that the drought sensitivity of Arabidopsis mutant in *ROOT HAIR DEFECTIVE 2* is linked to altered homeostasis of plasma membrane proteins and cytoskeleton remodelling.

## Introduction

NADPH oxidases, encoded in plants by *RESPIRATORY BURST OXIDASE HOMOLOG* (*RBOH*) genes, are plasma membrane (PM) localised proteins with pleiotropic functions in plant development and stress response. Ten *RBOH* isoforms have been identified in Arabidopsis (*AtRBOHA* to *AtRBOHJ*) with differing tissue-dependent and conditional expressions (Kaur and Pati, 2018; Chapman *et al*., 2019). Catalytic activity of NADPH oxidases leads to the formation of superoxide anion in the apoplast by transferring electrons from NAD(P)H to molecular oxygen. Under environmental stimuli, NADPH oxidases are activated by phosphorylation and Ca^2+^, leading to localised superoxide production. After the enzymatic or nonenzymatic conversion of superoxide to hydrogen peroxide, this is transported to intracellular space to convey the external signals (Chapman *et al*., 2019). To reveal a mode of RBOH regulation, it was shown that ROS production is also affected by Rho-of-Plant (ROP) GTPases. One of them, ROP6, is responsible for the formation of nanodomains under osmotic stress, where it recruits, interacts with, and activates RBOHD (Smokvarska *et al*., 2020). This RBOHD isoform undergoes nanodomain-associated endocytosis during salt stress (Hao *et al*., 2014).

Arabidopsis RBOHC/RHD2 (RHD2 stands for ROOT HAIR DEFECTIVE 2) represents an isoform preferentially expressed in roots and root hairs (Foreman *et al*., 2003). Root hair elongation is preconditioned by the polar accumulation of ROS produced by RBOHC and it cannot be replaced by RBOHJ or RBOHH (Kaya *et al*., 2019). Advanced quantitative microscopy revealed that GFP-RBOHC/RHD2 is delivered to the apical PM domain of the root hair by *trans*-Golgi network (TGN) in a structural sterol-dependent manner (Kuběnová *et al*., 2022). ROS production by RBOHC in root hairs depends on ROP2 (Jones *et al*., 2007) and SUPERCENTIPEDE (SCN1), a Rho GTPase GDP dissociaton inhibitor (Carol *et al*., 2005; Arenas-Alfonseca *et al*., 2018). Transcriptional regulation of *RBOHC* expression controlled by transcription factor RSL4 is essential for the root hair growth-promoting effect of auxin (Mangano *et al*., 2018). RBOHC is activated by CIPK26-mediated phosphorylation, but CIPK itself is unnecessary for root hair formation (Zhang *et al*., 2018). Moreover, the expression of *RBOHC* is stimulated by ethylene and is responsible for ROS production and root hair formation in response to ethylene (Martin *et al*., 2022).

Loss-of-function *root hair defective 2* (*rhd2*) mutants exert short, non-growing root hairs with missing tip-focused ROS and Ca^2+^ gradients (Schiefelbein and Somerville, 1990; Foreman *et al*., 2003) and this phenotype might be reversed by external addition of ROS or pH alkalinisation (Monshausen *et al*., 2007). The mutant also shows altered dynamics of early endosomes, while late endosomes appear unaffected (Kuběnová *et al*., 2022).

Although the mechanisms underlying defective root hair elongation in *rhd2-1* mutant are well known, physiological consequences at the level of shoots and whole plants have not been studied so far. Processes occurring downstream of altered ROS production and signalling in the *rhd2-1* mutant could be reflected by proteome composition of roots and shoots. To answer these questions, we performed a differential proteomic analysis on roots and shoots of *rhd2-1* mutant. Based on independent validations of differentially regulated proteins, we suggest that the loss of RBOHC function is associated with altered homeostasis of PM localised proteins, including aquaporin PIP1 isoform. This was accompanied by compromised drought stress tolerance in *rhd2-1* mutant.

## Material and Methods

### Plant Material and Growth Conditions

Seeds of *Arabidopsis thaliana*, ecotype Col-0, as a wild-type, *rhd2-1*, a loss-of-function mutant (Foreman *et al*., 2003), as well as transgenic Col-0 and *rhd2-1* plants bearing *35S::mRFP:TUB6* (mRFP-TUB6) construct (Fujita et al. 2011), as well as transgenic Col-0 and *rhd2-1* plants carrying *35S::GFP:FABD2* (GFP-FABD2) were used in this study. To prepare the *rhd2-1* transgenic lines with mRFP-TUB6 or GFP-FABD2, the *rhd2-1* mutant was crossed as a female donor with a Col-0 mRFP-TUB6 line or Col-0 GFP-FABD2 as male donors. For proteomic analysis and immunoblotting, seeds were ethanol-sterilised and grown for 14 days on ½ Murashige and Skoog (½ MS) media at 21°C, 70% humidity, and 16-h/8-h light/dark cycle. The illumination intensity was 130 µmol m^−2^.s^−1^. Roots and above-ground parts were harvested. The samples were harvested 6-8h after initiation of light illumination. Three days-old seedlings were used for live-cell imaging and whole-mount immunolocalization. To test the response of plants to drought stress, seeds were placed to substrate in the pots and the plants were grown for 4 weeks under the conditions mentioned above.

### Proteomic Analysis

Peptide digests were prepared as described previously (Takáč *et al*., 2017). Briefly, the harvested biological material was grounded to a fine powder in liquid nitrogen, and the proteins were extracted by phenol extraction and methanol precipitation. Proteins were digested in solution by sequencing-grade modified trypsin. Proteins identification, data filtering, quantification and statistical evaluation were performed as described in Melicher *et al*. (2022). The LC-ESI-MS/MS analysis was performed on a nanoflow HPLC system (Easy-nLC1200, Thermo Fisher Scientific) coupled to an Orbitrap Fusion Lumos mass spectrometer (Thermo Fisher Scientific, Bremen, Germany) equipped with a nano-electrospray ionisation source. Peptides were first loaded onto a trapping column and subsequently separated on a 15 cm C18 column (75 μm × 15 cm, ReproSil-Pur 5 μm 200 Å C18-AQ, Dr. Maisch HPLC GmbH, Ammerbuch-Entringen, Germany). The mobile phase consisted of water with 0.1% formic acid (solvent A) and acetonitrile/water (80:20 (v/v)) with 0.1% formic acid (solvent B). Peptides were eluted with a linear 110 min gradient from 5% to 21% solvent B in 62 min and then to 36% solvent B in 110 min, followed by a wash stage with 100% solvent B. MS data were acquired automatically using Thermo Xcalibur 4.1. software (Thermo Fisher Scientific). An information-dependent acquisition method consisted of an Orbitrap MS survey scan with a mass range of 300–1750 m/z followed by HCD fragmentation for the most intense peptide ions.

Data files were searched for protein identification using Proteome Discoverer 2.3 software (Thermo Fisher Scientific) connected to an in-house server running the Mascot 2.7.0 software (Matrix Science). Data were searched against the SwissProt database (version 2019_11) using the *A. thaliana* taxonomy filter. The following parameters were used: static modifications: carbamidomethyl (C)*, variable modifications: oxidation (M), acetyl (protein N-term), peptide mass tolerance: ± 10 ppm, fragment mass tolerance: ± 0.02 Da, maximum missed cleavages: 2, FDR = 0.01. Methionine oxidation is a common modification during sample processing and is generally included in the search parameters. Only unique peptides were used for the protein level quantifications. Abundance values for peptides and proteins were calculated based on the intensities of peptide precursor ions. All peptides were used for normalisation. T-test (background based) was used as a quantification hypothesis test. Minimum two replicates in one sample group had to contain the same quantification feature in order to be included into the quantification. Proteomics analyses were carried out with three biological replicates for each of the four biological samples (roots and aerial parts of Col-0 and *rhd2-1*, respectively). Each replicate contained at least 30 seedlings. All peptides were used for normalisation. ANOVA (adjusted p ≤ 0.05) was used to filter statistically significant results, applied to proteins exhibiting a fold change ≥ 1.5. Proteins identified by one peptide were excluded from the analysis. Proteins present in all three replicates corresponding to the control proteome and absent in all three replicates of the test proteome were considered unique for the control proteome and vice versa.

### Bioinformatics analysis of the differential proteome

The differential proteome was evaluated using GO Term Enrichment tool on https://www.arabidopsis.org/tools/go_term_enrichment.jsp. Protein localisation was predicted using WoLF PSORT (https://wolfpsort.hgc.jp/). For analysis of protein interaction networks representation in differential proteome, STRING (Szklarczyk *et al*., 2019) web-based application was used considering experimental evidences and co-expression while the minimum required interaction score was set to 0.7 (relevant for high confidence prediction).

### Immunoblotting Analysis

Protein extraction and immunoblotting analysis were performed as described previously (Melicher *et al*., 2022). To detect abundances of PIPs and CHC, anti-PIP1.1-5 (AS09 487) and anti-CHC antibodies (AS10 690; all from Agrisera, Sweden) were used, respectively. Band densities were quantified using Image Lab software (Bio-Rad). Three biological replicates were carried out in this analysis and differences between the Col-0 and *rhd2-1* mutant were evaluated by one-way ANOVA analysis with post hoc Tukey HSD test. To test the loading of equal amount of proteins, proteins were visualised on membranes using Ponceau-S staining.

### Whole Mount Immunolocalization

Three days-old seedlings of Col-0 and *rhd2-1* were used for immunolocalization of PIP1 in roots as described previously (Šamajová *et al*., 2014). In the case of leaf immunolabeling, we used leaves of 10 days-old Col-0 and *rhd2-1* plants and followed the protocol by Pasternak *et al*. (2015). Samples were immunolabeled with anti-PIP1.1-5 rabbit polyclonal primary antibody (Agrisera, Sweden) diluted 1:150 in 3% (w/v) BSA in PBS at 4°C overnight. Then, AlexaFluor 647 goat anti-rabbit secondary antibody (Abcam) diluted 1:500 in 2.5% (w/v) BSA in PBS was used for 2 h incubation at 37°C. Microscopic analysis of immunolabeled samples was performed with a Zeiss 880 LSM Airyscan equipped with 32 GaAsP detector (Carl Zeiss, Germany) using a Plan-Apochromat 63×/1.4 Oil DIC M27 objective and fluorophore excitation with 631 nm laser. The image acquisition, post-processing and fluorescence intensity measurement were made using Zeiss ZEN software (Black and Blue versions, Carl Zeiss, Germany). Semiquantitative evaluation of immunolabeled PIP1 was performed using fluorescence intensity profile measurement of PIP1 spot-like structures localised at the PM of root and leaf epidermal cells. Data were obtained from root and leaf epidermal cells (40 cells per root; 15 cells per leaf) from 3 individual plants of Col-0 and *rhd2-1* mutant. Statistical significance was evaluated using t-test (**p < 0.01).

### Drought stress

Col-0 and *rhd2-1* plants grown in pots for 4 weeks as described above, were subjected to drought stress by withholding the irrigation, while in controls, plants were continuously irrigated. Following 18 days, plants were documented using a Nikon 5000 SLR camera.

### In vivo observation of microtubules and actin using lattice light-sheet fluorescence microscopy

The study of cortical microtubules in transgenic Col-0 and *rhd2-1* lines carrying mRFP-TUBULIN β6 fluorescent marker (Col-0 mRFP-TUB6 and *rhd2-1* mRFP-TUB6, respectively) in root cells was performed by lattice light-sheet fluorescence microscopy (lattice LSFM, Chen *et al*., 2014). High-resolution single-side illumination and single-side detection lattice LSFM system was equipped with illumination objective 28.5×/0.62 NA, detection objective 25×/1.1 NA, and Orca Flash 4.0 sCMOS camera. Exposure time for image acquisition was 100 ms. Fluorescence of mRFP was excited at 589 nm. Images were acquired at 1 s intervals for 60 time points at Z-stack mode containing 20 optical sections with the step size of 0.25 μm. The Richardson-Lucy algorithm was used for deconvolution. Images are presented as orthogonal projections of individual planes.

Orientation of cortical microtubules in root epidermal cells of Col-0 mRFP-TUB6 and *rhd2-1* mRFP-TUB6 plants, with respect to the orientation of root longitudinal axis, was analysed using the OrientationJ plugin in Fiji (Schindelin *et al*., 2012; http://sites.imagej.net/BIG-EPFL/). Mean values of microtubule orientation, their analysis and comparisons between lines over time of recordings were evaluated, and the Python programming language with Anaconda Navigator program, version 2.1.4 (Anaconda, Inc.) was used for data graphical presentation. The values with a low coherence (0.02 and below) were excluded from the analysis. Graphical presentations of cortical microtubule distribution were prepared in Microsoft Excel.

### Lattice LSFM data processing using PIV analyser

Velocity of actin filaments in epidermal root cells of Col-0 GFP-FABD2 and *rhd2-1* GFP-FABD2 transformed plants was analysed using the PIV analyser plugin in Fiji (https://imagej.net/plugins/piv-analyser). Mean values of actin velocities, their analysis, and comparisons between lines over time of recordings were evaluated, and a Python programming language with Anaconda Navigator program, version 2.1.4 (Anaconda, Inc.) was used for data graphical presentation.

## Results

### The proteome of Arabidopsis rhd2-1 mutant substantially differs from the Col-0

Altogether, 121 and 108 differentially regulated proteins were found in *rhd2-1* roots and above-ground parts, respectively, compared to the Col-0 (Supplementary Tables S1 and S2). Gene ontology (GO) annotation analysis of the differential proteome showed that the highest number of differentially regulated proteins is involved in metabolism, response to abiotic and biotic stimulus, development, gene expression and translation. The *rhd2-1* also differed from the Col-0 in proteins involved in photosynthesis, intracellular transport, response to hormones, lipid, and cell wall modification (Supplementary Table S3). The annotations such as metabolism, gene expression (roots), translation (both samples), starch metabolic process (above-ground parts) and response to gravity (roots) were enriched in differential proteomes, as evaluated by Fisher’s exact test (Supplementary Figures S1, S2; Supplementary Table S3). Proteins involved in translation formed the biggest protein interaction cluster in the differential proteomes (Figure 1). Most differentially regulated proteins localise to the cytoplasm, nuclei, plastids, vacuoles, PM, cell cortex, ribosome, plasmodesmata, cell junctions, symplast and vesicles. Annotations such as ribosome, vacuolar membrane, vacuole, plasmodesma, nucleolus, cell wall, cytosol and cytoplasmic vesicle were enriched in the differential proteomes (Supplementary Figures S3, S4; Supplementary Table S3).

**Figure 1.**
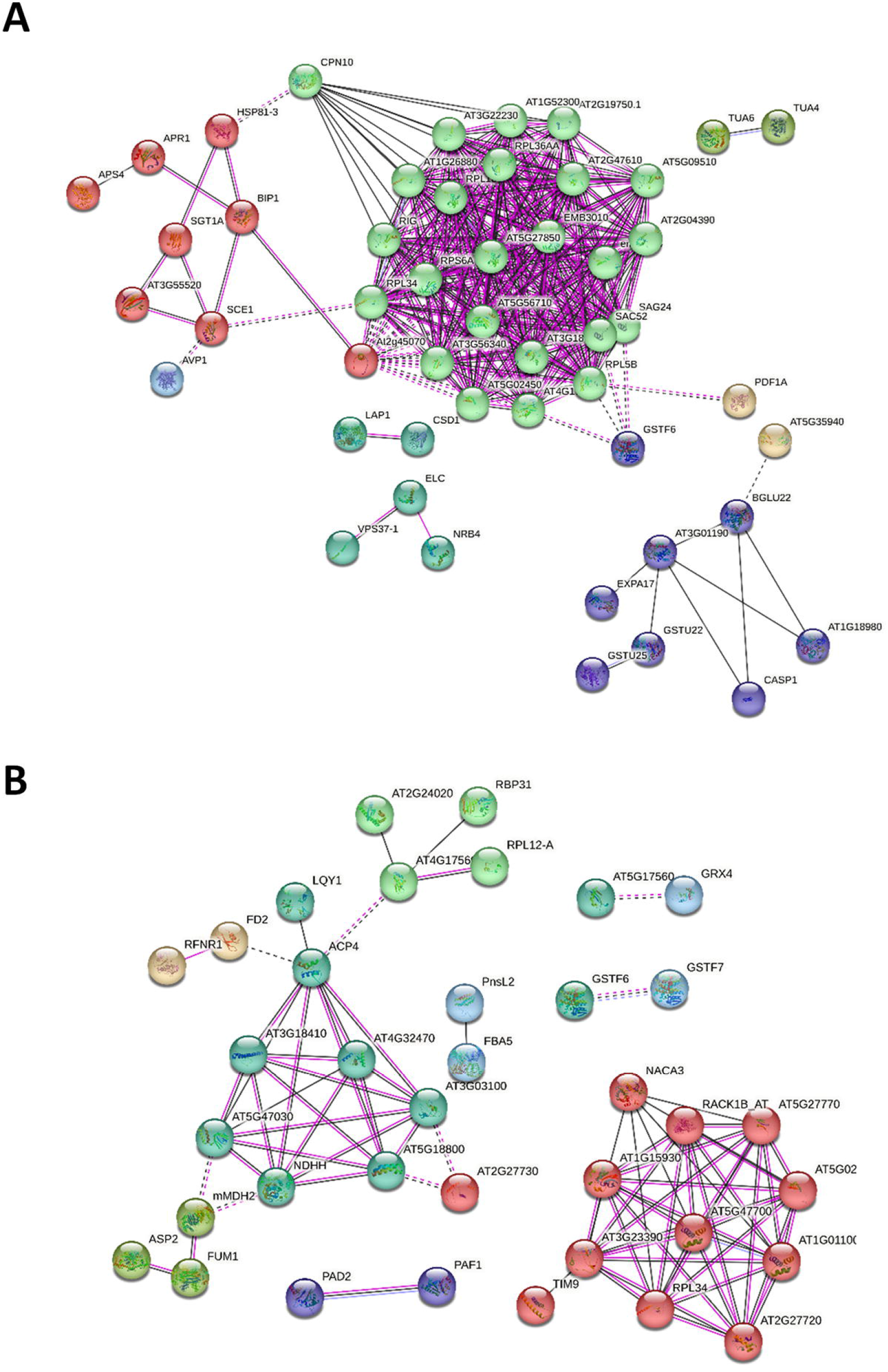
Presentation of protein interaction networks in the differential proteome of roots (A) and above-ground parts (B) of Arabidopsis *rhd2-1* mutant as compared to Col-0. The nodes represent individual proteins. Only experimentally proved interactions (also including heterologous proteins) and co-expressions are considered. The node colours indicate common Kmeans clustering, as evaluated by STRING. Prevalent functional annotations in clusters in A: light green – ribosomal proteins; red - protein folding and degradation; purple – proteasome complex; green - tubulin isoforms; dark blue - cell wall modification and glutathione S-transferases; dark green - membrane trafficking, Cu/Zn superoxide dismutase 1, cytosol aminopeptidase. B:turquoise – mitochondrial electron transport chain, glutathione S-transferases, glutaredoxin; green - TCA cycle enzymes; light brown – ferredoxin and ferredoxin NADP reductase; light green – chloroplastic ribosomal proteins; red – ribosomal proteins; purple – proteasome subunits

Remarkably, differential proteomes of roots differed from above-ground parts mainly in defence-related proteins, glutathione S-transferases (GSTs), cell wall regulatory proteins and secreted proteins (Supplementary Figure S5). In roots of *rhd2-1* compared to Col-0, all defence-related proteins had a higher abundance, while in above-ground parts, they mainly showed a lower abundance. Also, secreted proteins and proteins involved in cell wall modification were much more affected in roots compared to above-ground parts.

In order to estimate which of the differentially regulated proteins contribute to the root hair phenotype of the mutant, we used the ePlant gene expression database (Fucile *et al*., 2011) to screen the genes encoding the differentially regulated proteins expressed in trichoblast cells, or in the root differentiation zone. Proteins involved in membrane trafficking and metabolism had root hair-, or root differential zone-specific expression patterns (Table 1). Remarkably, the majority of membrane trafficking proteins showed lower abundance in the *rhd2-1* mutant proteome, implying that these might contribute to its root hair phenotype. Other proteins included PEROXIDASE 27, CRYPTOCHROME DASH, a protein involved in vacuolar transport MONOSACCHARIDE-SENSING PROTEIN 1 and proteins involved in signalling (CSC1-LIKE PROTEIN ERD4, PHOSPHOINOSITIDE PHOSPHATASE SAC8; Table 1).

**Table 1.** Proteins differentially regulated in roots of *rhd2-1* mutant potentially contributing to its root hair phenotype. Proteins were selected considering prevalent expression of their encoding genes in trichoblasts, or in the root differentiation zone with root hairs (ePlant; https://bar.utoronto.ca/eplant/)

| Accession | Localisation | Function | Protein name | Fold change |
| --- | --- | --- | --- | --- |
| Q9LFW1 | cytoplasm, cytosol, Golgi apparatus | cell wall | UDP-arabinopyranose mutase 2 | 1.59 |
| F4IVL6 | endosomal membrane | membrane trafficking | DnaJ homolog subfamily C GRV2 | 0.01 |
| P38389 | ER membrane; single-pass membrane protein | membrane trafficking | Protein transport protein Sec61 subunit beta | 0.62 |
| Q9FFF3 | Golgi apparatus membrane | membrane trafficking | Conserved oligomeric Golgi complex subunit 1 | 0.60 |
| F4IXW2 | cytoplasm, cytosol, Golgi apparatus, trans-Golgi network membrane | membrane trafficking | Brefeldin A-inhibited guanine nucleotide-exchange protein 5 | 0.62 |
| O82629 | vacuolar membrane | membrane trafficking | V-type proton ATPase subunit G2 | 0.49 |
| Q39055 | mitochondrion matrix | metabolism | GTP 3',8-cyclase, mitochondrial | 0.37 |
| Q8LDK3 | mitochondrion inner membrane | metabolism | NADH dehydrogenase [ubiquinone] 1 beta subcomplex subunit 2 | 0.55 |
| Q96328 | ER membrane | phospholipid signalling | Phosphoinositide phosphatase SAC8 | 0.46 |
| Q84KJ5 | plastid, chloroplast, mitochondrion | photoreceptor | Cryptochrome DASH, chloroplastic/mitochondrial | 0.01 |
| Q43735 | secreted | redox | Peroxidase 27 | 0.54 |
| Q9C8G5 | membrane | signalling | CSC1-like protein ERD4 | 0.53 |
| Q96290 | membrane | transport | Monosaccharide-sensing protein 1 | 0.57 |

### Deregulation of stress-related proteins in the rhd2-1 mutant

Differences accounted between the *rhd2-1* and Col-0 occurred in proteins involved in plant responses to drought, high temperature, xenobiotics and biotic stress. The *rhd2-1* mutant showed downregulation of aquaporins PIP1-1 in roots, while PIP1-3 was upregulated in above-ground parts (Supplementary Figure S5). Immunoblotting analysis using a primary antibody recognising five PIP1 isoforms (PIP1-1 to PIP1-5) validated the proteomic analysis, showing that protein abundance corresponding to PIP1 isoforms was lower in the *rhd2-1* roots, but higher in the above-ground plant parts (Figure 2A-D). These results were further corroborated by immunolabeling of *rhd2-1* and Col-0 roots and leaves using the above-mentioned antibody (Figure 2E-H). Semiquantitative fluorescence intensity profile measurement of spot-like structures of PIP1 isoforms localised in the PM of *rhd2-1* root epidermal cells showed decreased intensities compared to Col-0 (Figure 2E-F). However, increased fluorescence intensity of PIP1 isoforms localised in the PM was observed in leaf epidermal cells of the *rhd2-1* mutant (Figure 2G-H). These results are consistent with proteomic and immunoblotting analyses. The differential abundance of a water channel aquaporins may indicate the altered response of the *rhd2-1* mutant to drought stress. In addition, other 30 differentially abundant proteins (15 in roots and 15 in above-ground parts), encoded by genes showing increased expression in response to drought (ePlant), were downregulated in *rhd2-1* mutant (Figure 3). These include proteins involved mainly in abiotic stress response, membrane trafficking, metabolism, signalling and transmembrane transport. All these data might point to altered drought stress tolerance of *rhd2-1* mutant. To test this hypothesis, we examined the drought stress tolerance of Col-0 and *rhd2-1* plants growing in soil by withholding their irrigation. This analysis revealed a higher susceptibility of *rhd2-1* compared to Col-0 (Figure 4).

**Figure 2.**
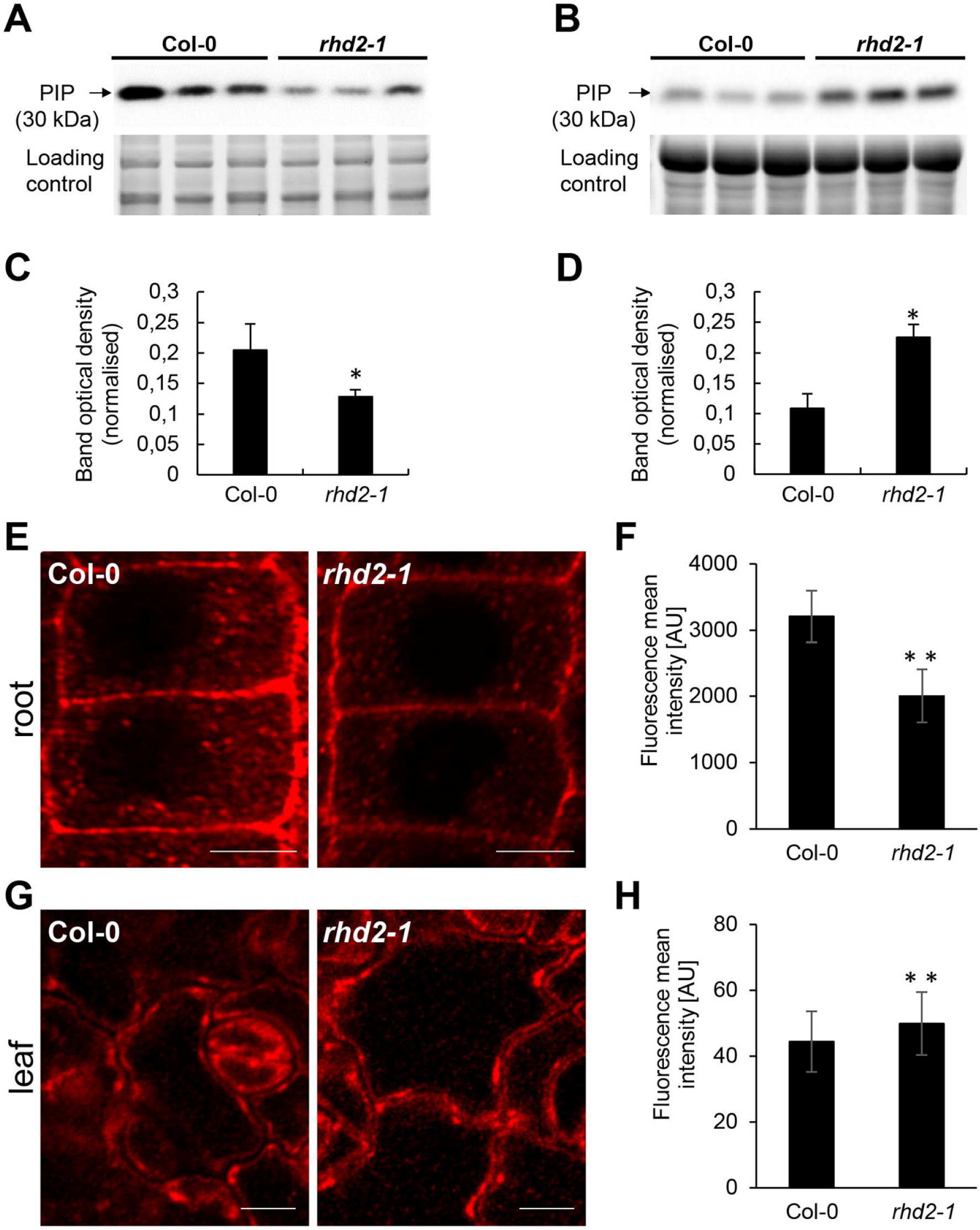
Immunoblotting abundance analysis and immunolocalization of PLASMA MEMBRANE INTRINSIC PROTEIN 1 (PIP1) in Col-0 plants and *rhd2-1* mutant. Immunoblots of PIP1 in root system (A) and above-ground part (B) in Col-0 and *rhd2-1*. Each of the three lanes within one genotype represents a biological replicate. (C, D) Quantification of band optical densities in A and B. Immunoblots are supplemented with respective controls of protein loading using Stain-free gels. Values are expressed as mean ± SD (N = 3) and asterisks indicate a statistically significant difference as revealed by one-way ANOVA with post-hoc Tukey HSD test (p < 0.05). (E, G) Subcellular immunolocalization of PIP1 in Col-0 and *rhd2-1* mutant root (E) and leaf (G) epidermal cells. (F, H) Semiquantitative analysis of PIP1 spot-like structures fluorescence intensity on the plasma membrane of root and leaf epidermal cells. Data are obtained from root and leaf epidermal cells (40 cells per root; 15 cells per leaf) from 3 individual plants of Col-0 and *rhd2-1* mutant. Statistical significance was evaluated using t-test (**p < 0.01). Asterisks indicate statistical significance between Col-0 and *rhd2-1* mutant (**p < 0.01). Error bars show ± standard deviation (SD). Both analyses were performed using anti-PIP1.1-5 antibody. Bar = 5 µm.

**Figure 3.**
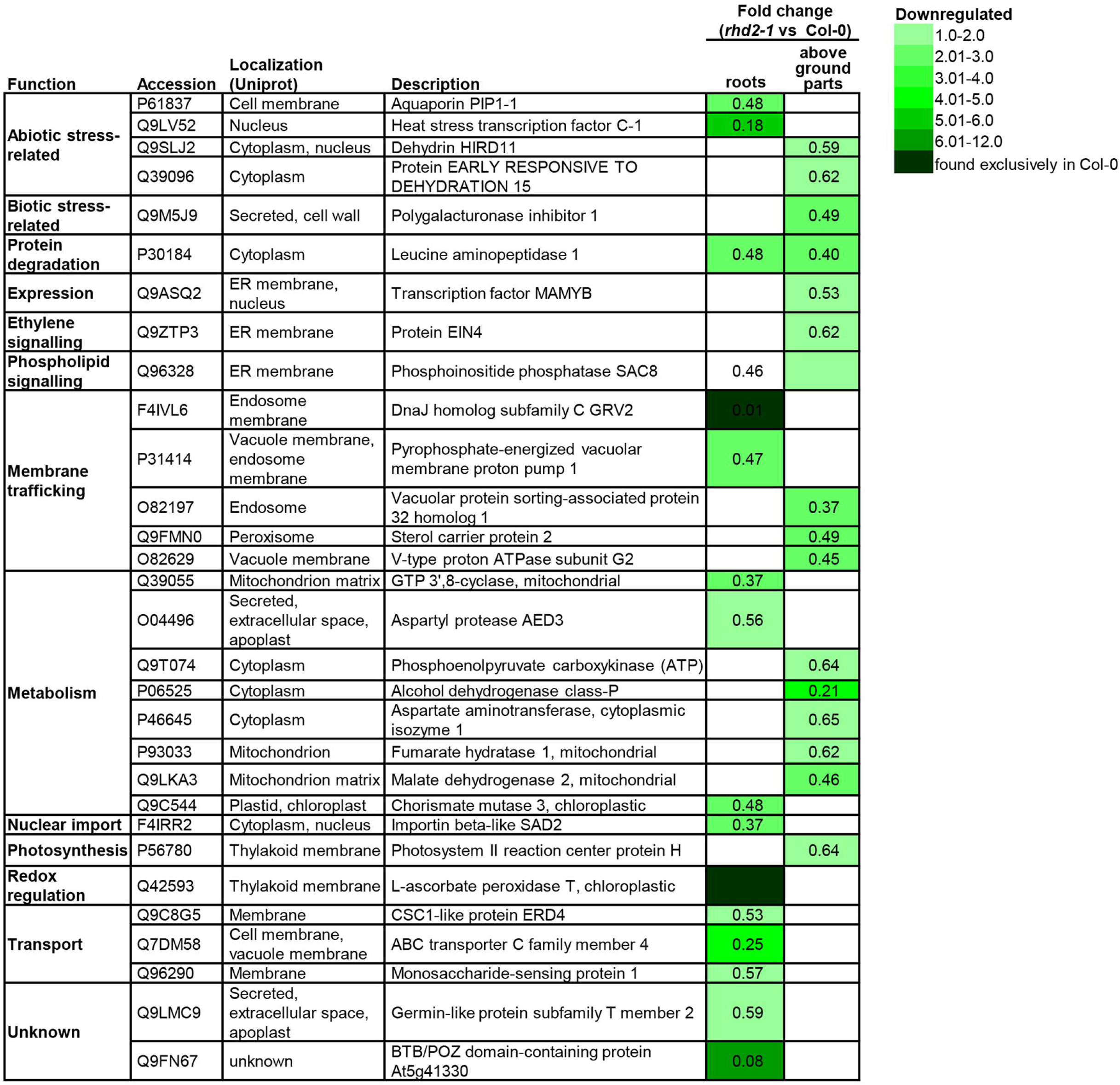
Heat map showing the abundance ratio of proteins differentially regulated in *rhd2-1* mutant and Col-0, likely contributing to the altered response of *rhd2-1* mutant to drought stress. The expression of genes encoding the listed proteins were found as upregulated after drought stress in ePlant expression database. Fold changes are presented in colour as indicated by the colour key on the right. Proteins unique for either control or treated samples were found solely in the proteomes of control or treated samples, considering all analysed replicates.

**Figure 4.**
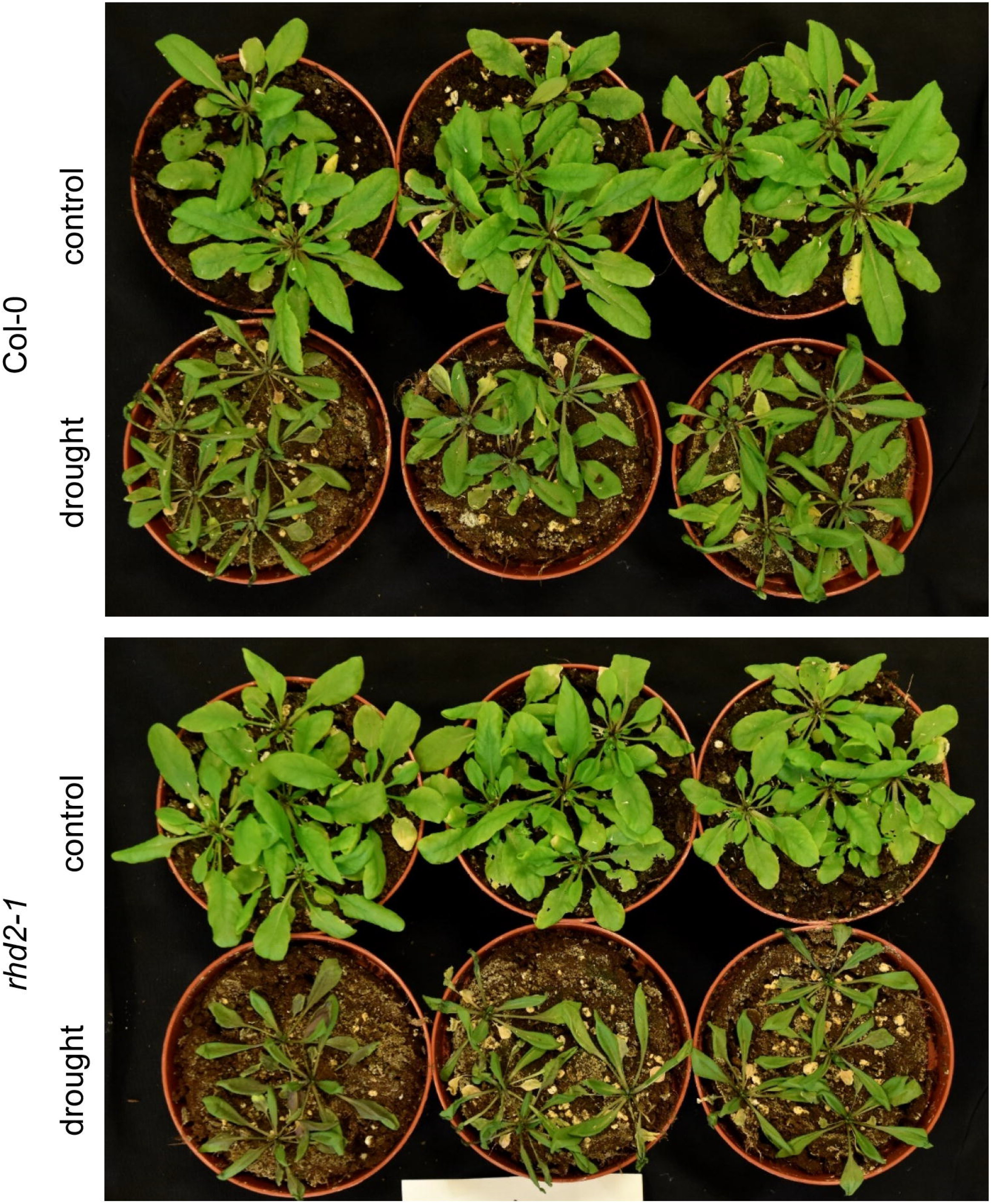
Effects of drought stress on development and resistance of *rhd2-1* mutants and Col-0 plants in soil. Note a higher sensitivity of mutant to drought stress.

The *rhd2-1* mutant also showed prevalently lower abundances of proteins involved in heat stress response, including HEAT STRESS TRANSCRIPTION FACTOR C-1, HSP90-3 and HEAT SHOCK FACTOR-BINDING PROTEIN. On the other hand, heat shock 70 kDa protein BIP1 showed higher abundance in *rhd2-1* roots (Supplementary Tables S1 and S2).

GLUTATHIONE S-TRANSFERASES (GSTs) are responsible for conjugating diverse xenobiotics to glutathione. The differential proteome of *rhd2-1* contained seven isoforms of GSTs, including GSTU5 (roots), GSTU22 (both plant parts), GSTU25 (roots), GSTL1 (roots), GSTF2 (above-ground parts), GSTF6 (both plant parts) and GSTF7 (above-ground parts; Supplementary Tables S1 and S2). While all isoforms found in roots had considerably increased abundance in the *rhd2-1* mutant, three out of four isoforms were downregulated in above-ground parts (Supplementary Figure S5). Glutathione-xenobiotics complexes are transported to the vacuole by ABC transporters (Chronopoulou *et al*., 2017). Two candidates for these transporters were found in our proteomic screens. ABC TRANSPORTER C FAMILY MEMBER 4 showed downregulation in *rhd2-1* roots, while ABC TRANSPORTER G FAMILY MEMBER 32 was upregulated in the above-ground parts (Supplementary Figure S5).

Furthermore, *RHD2* functional mutation leads to changes in the abundance of proteins involved in biotic stress response. Two important proteins involved in the perception of pathogen-associated molecular patterns (PAMPs), serine/threonine-protein kinase BSK1 and MIK2 receptor kinase were upregulated in *rhd2-1* roots. Some other proteins, although not being a part of PAMP signalling but known to be involved in plant defense response, were also upregulated (Supplementary Figure S5). On the other hand, similar defense-related proteins were downregulated in above-ground parts of *rhd2-1* mutant (Supplementary Figure S5).

Proteins involved in antioxidant defense and redox regulation are also deregulated in the *rhd2-1* mutant. Two THIOREDOXIN isoforms and one GLUTAREDOXIN were downregulated, but proteins involved in the synthesis of nonenzymatic antioxidants were upregulated in above-ground parts of *rhd2-1*. Roots of *rhd2-1* exhibited an increased abundance of enzymatic antioxidants, such as Cu Zn SUPEROXIDE DISMUTASE 1 (CSD1) and MONODEHYDROASCORBATE REDUCTASE 3 (MDHAR3), however, a thylakoid ASCORBATE PEROXIDASE (tAPX) was downregulated (Supplementary Tables S1 and S2).

### RHD2 mutation leads to altered homeostasis of plasma membrane, tonoplast, vesicular trafficking and cytoskeletal proteins

As annotated by Uniprot, the differential proteome of *rhd2-1* mutant contained twelve proteins localised in the PM and four in the tonoplast. Proteome of *rhd2-1* differed in the abundance of PM localised proton-, cutin- and multidrug-transporting proteins and two aquaporin isoforms PIP1-1 and PIP1-3 (mentioned above), localised in the PM. Moreover, proteins involved in Casparian strip and cellulose microfibril formation, intercellular movement across plasmodesmata and perception of microbe-associated molecular patterns (MAMPs) showed differential abundance between the mutant and Col-0 (Figure 5).

**Figure 5.**
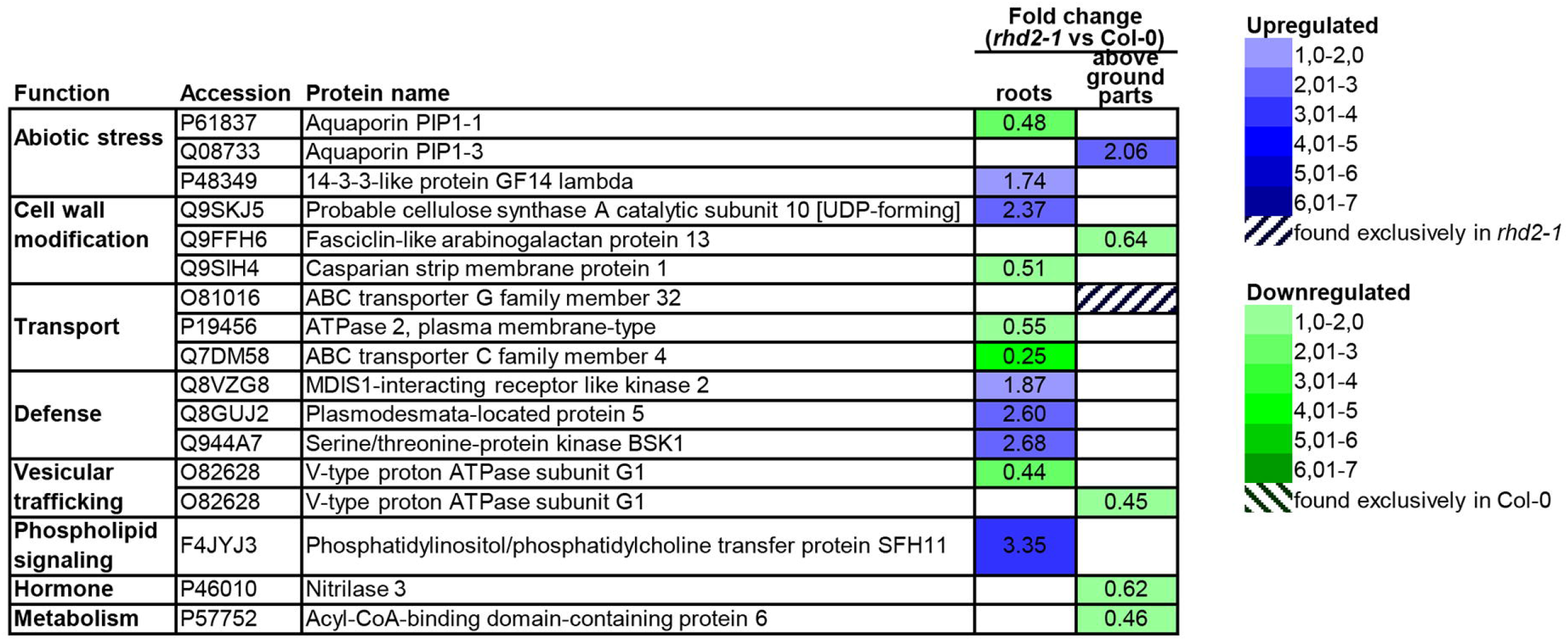
Heat map showing the abundance ratio of proteins localised to the plasma membrane and differentially regulated in the *rhd2-1* mutant versus Col-0. Fold changes are presented in colour as indicated by the colour key on the right. Proteins unique for either control or treated samples were found solely in the proteomes of control or treated samples, considering all analysed replicates.

Pumps transporting protons across tonoplast, such as PYROPHOSPHATE-ENERGIZED VACUOLAR MEMBRANE PROTON PUMP 1, and two subunits of V-TYPE PROTON ATPase (G1 and G2) were downregulated in the *rhd2-1* mutant. MONOSACCHARIDE-SENSING PROTEIN 1, a protein transporting monosaccharides to the vacuole, was also downregulated. TONOPLAST SLOCALISED HEVEIN-LIKE PREPROTEIN, a pathogenesis-related protein, was also affected (Figure 5).

A considerable number of proteins is predicted to be secreted to the extracellular space. They include cell wall constituents, proteins involved in cell wall modification, proteases and lectins. Notably, cell wall modifying enzymes were much more abundant in roots compared to above-ground parts in *rhd2-1* (Supplementary Figure S5).

Our proteomic data point also to malfunction of vesicular transport in *rhd2-1* mutant due to the changed abundance of 3 components of ESCRT endosomal sorting complex, a component of the retromer complex, two components of SNARE (SOLUBLE N-ETHYLMALEIMIDE SENSITIVE FACTOR ATTACHMENT PROTEIN) complex, a subunit of conserved oligomeric Golgi complex, and exocyst complex (Table 2). In addition, CLATHRIN HEAVY CHAIN 1 was 1.78 fold upregulated in above-ground parts of *rhd2-1* (Table 2). Identified proteins with altered abundances leading to malfunction of vesicular trafficking in *rhd2-1* mutant may directly cause misbalance in PIPs distribution and abundance in membranes, as vesicles are responsible for their targeted delivery. Independent validation by immunoblotting analysis supported upregulation of CLATHRIN HEAVY CHAIN in above-ground parts of *rhd2-1* (Figure 6), in agreement with the proteomic data.

**Figure 6.**
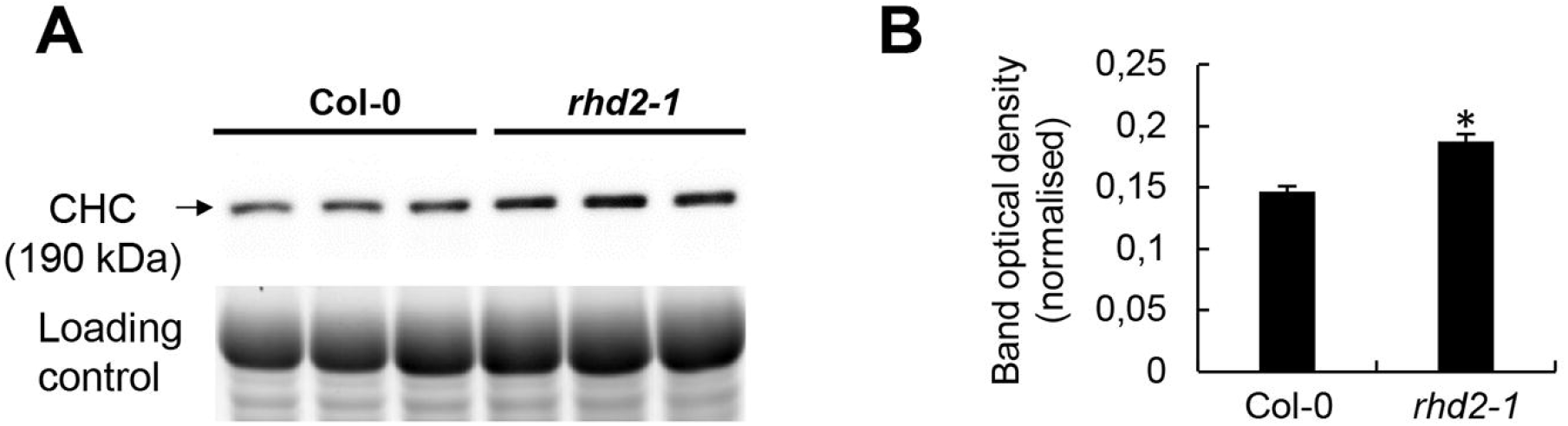
Immunoblotting analysis of CLATHRIN HEAVY CHAIN (CHC) in Col-0 and *rhd2-1* plants. (A) Immunoblots of CHC in above-ground part of Col-0 and *rhd2-1*. (B) Quantification of band optical densities in A. The immunoblot is supplemented with respective controls of protein loading using Stain-free gels. Values are expressed as comparison of Col-0 to *rhd2-1* (mean ± SD, N = 3) and asterisks indicate a statistically significant difference as revealed by one-way ANOVA with post-hoc Tukey HSD test (p < 0.05).

**Table 2.** Differentially abundant vesicular transport proteins in roots and above-ground parts of *rhd2-1* mutant.

| Accession | Protein name | Fold change<br>(*rhb2-1*<br>vs Col-0) | Molecular function,<br>complex |
| --- | --- | --- | --- |
| **Root differential proteome** |  |  |  |
| O82628 | V-type proton ATPase subunit G1 | 0.441 | vacuolar protom pump |
| O82629 | V-type proton ATPase subunit G2 | 0.488 | vacuolar protom pump |
| P31414 | Pyrophosphate-energized vacuolar membrane proton pump 1 | 0.47 | vacuolar protom pump |
| F4IXW2 | Brefeldin A-inhibited guanine nucleotide-exchange protein 5 (BIG5) | 0.62 | ADP-ribosylation factor (ARF) guanine nucleotide exchange factor |
| Q9STT2 | Vacuolar protein sorting-associated protein 29 | 1.52 | Retromer complex subunit |
| Q9LHG8 | Protein ELC (VPS23) | 2.98 | ESCRT complex subunit |
| Q9SCP9 | Vacuolar protein-sorting-associated protein 37 homolog 1 | 2.08 | ESCRT complex subunit |
| P93654 | Syntaxin-22 | 0.57 | SNARE complex |
| Q9SY60 | Exocyst complex component EXO84C | 2.34 | Exocyst complex |
| Q9FFF3 | Conserved oligomeric Golgi complex subunit 1 | 0.60 | Conserved oligomeric Golgi complex subunit |
| P38389 | Protein transport protein Sec61 subunit beta | 0.62 | unknown |
| Q9ZPS7 | Transmembrane 9 superfamily member 3 | 0.34 | unknown |
| F4IVL6 | DnaJ homolog subfamily C GRV2 | 0.01 | unknown |
| **Above-ground plant part differential proteome** |  |  |  |
| O82628 | V-type proton ATPase subunit G1 | 0.45 | vacuolar protom pump |
| O82629 | V-type proton ATPase subunit G2 | 0.45 | vacuolar protom pump |
| Q0WNJ6 | Clathrin heavy chain 1 | 1.78 | clathrin coat |
| O82197 | Vacuolar protein sorting-associated protein 32 homolog 1 | 0.37 | ESCRT complex subunit |
| Q944A9 | Novel plant SNARE 11 | unique in *rhb2-1* | SNARE complex |

Two alpha TUBULIN isoforms (TUA2 and TUA6) showed higher abundances in the *rhd2-1* mutant compared to Col-0 (Supplementary Tables S1 and S2). These data, together with the simultaneous upregulation of MICROTUBULE ASSOCIATED PROTEIN 70-2 (MAP70-2) indicate possible new link between NADPH oxidases and microtubule organisation. This finding may pose a high significance to other processes in *rhd2-1* mutant to be affected at the proteome level, like altered abundances of PIPs, which are eventually transported and targeted to PM (similarly to cellulose synthase complexes) using cortical microtubules. Therefore, we generated transgenic Col-0 and *rhd2-1* lines expressing a microtubule marker mRFP-TUB6 and performed a detailed advanced lattice LSFM analysis to reveal the cortical microtubule organisation in root epidermal cells. We did not observe obvious differences between Col-0 and *rhd2-1* in the microtubule organisation in epidermal cells in the transition zone of the primary root (Figure 7A, B, E). Microtubules were oriented prevalently perpendicular to the root longitudinal axis (Figure 7A, B) with negligible deviations in microtubule orientation angles (Figure 7E). On the other hand, microtubules in epidermal cells of the root elongation zone showed higher level of anisotropy in the *rhd2-1* mutant compared to Col-0 (Figure 7C, D) with shift in the orientation angles (Figure 7F). Dynamic time-lapse lattice LSFM imaging of cortical microtubule orientation revealed similar trend between root epidermal cells of Col-0 and *rhd2-1* in the root transition zone (Figure 7G), but substantially different trend in the root elongation zone (Figure 7H).

**Figure 7.**
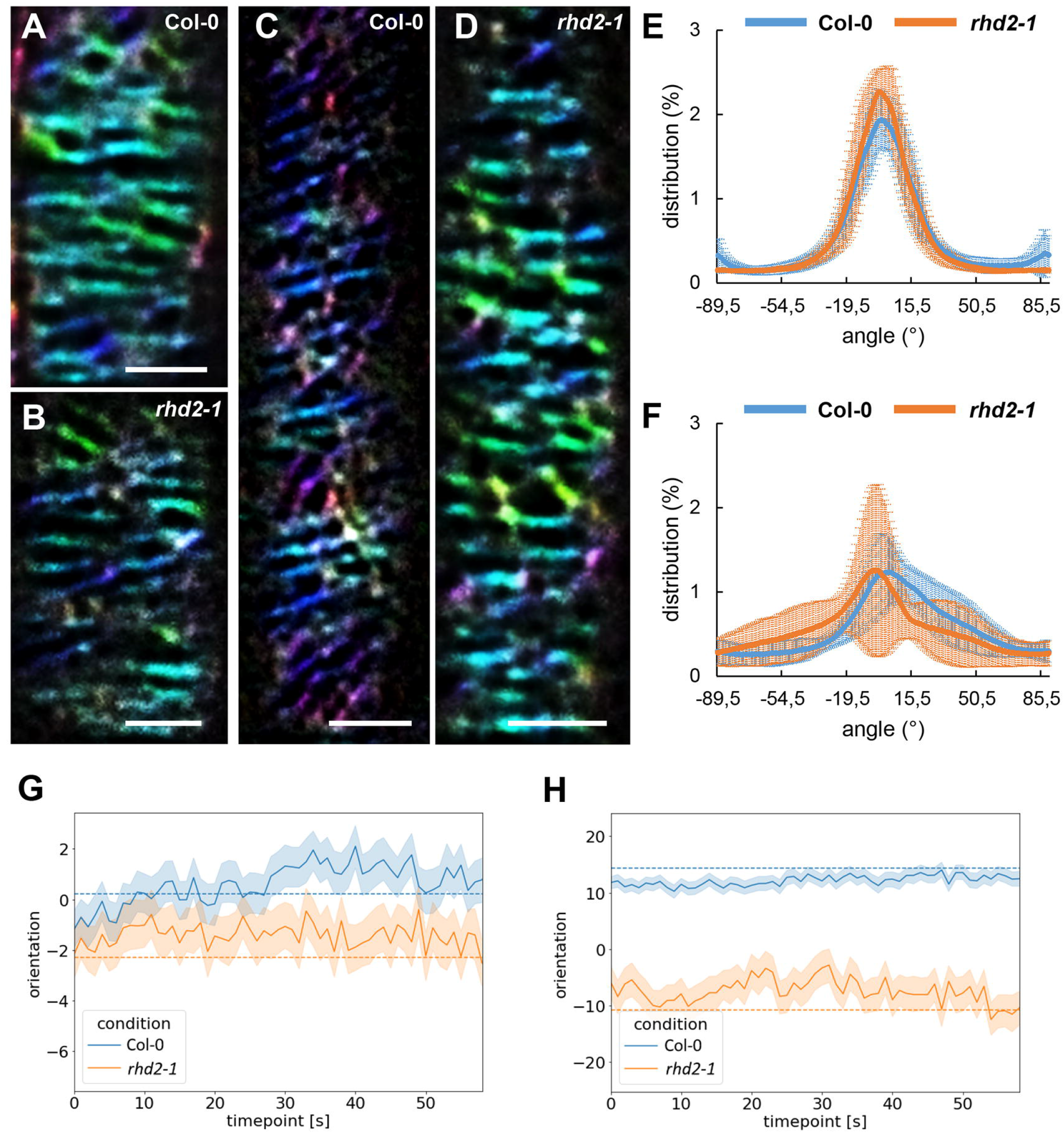
Analysis of cortical microtubule orientation in root epidermal cells of Col-0 and *rhd2-1* using lattice light-sheet fluorescence microscopy. (A-D) Color-coded presentation of cortical microtubule orientation in regard of the root longitudinal axis visualised by OrientationJ plugin of the Fiji. (A-B) Cortical microtubules in root epidermal cells from the transition zone of the Col-0 mRFP-TUB6 (A) and *rhd2-1* mRFP-TUB6 transgenic lines (B). (C-D) Cortical microtubules in root epidermal cells from the elongation zone of the Col-0 mRFP-TUB6 (C) and *rhd2-1* mRFP-TUB6 transgenic lines (D). (E-F) Distribution graphs of the cortical microtubule orientation from cells of the transition zone (E) and cells of the elongation zone (F). (G-H) Mean values of cortical microtubule orientation from 60 s long recording in cells of the transition zone (G) and cells of the elongation zone (H). Measurements are from 10 cells and 7 plants in Col-0, and from 5 cells and 3 plants in *rhd2-1* mutant in the transition zone, and from 7 cells and 5 plants in Col-0, and from 3 cells and 3 plants in *rhd2-1* mutant in the elongation zone. Bar = 5 µm.

Due to the substantial deregulation of proteins involved in membrane trafficking and the known dependence of the vesicular trafficking on actin cytoskeleton dynamics, we also aimed to examine the actin cytoskeleton in roots of *rhd2-1* mutant. The hypothesis indicating a possible alteration of actin cytoskeleton in *rhd2-1* mutant is supported also by decreased expression of genes encoding actin-binding proteins in *rhd2-1* transcriptome (Jones et al., 2006). We analysed actin velocity in root epidermal cells of Col-0 GFP-FABD2 and *rhd2-1* GFP-FABD2 transgenic lines using lattice LSFM. Velocity of actin filaments was semi-quantitatively analysed by colour-coding determination of movement speed in nm/s, supplemented by vectoral display of changes. This analysis combined velocity fluctuations with their spatial distribution, characterising thus actin filament dynamics in more thorough manner (Figure 8). It revealed a considerable reduction of actin cytoskeleton dynamics in *rhd2-1* mutant (Figure 8A-D). Direct time-lapse recording of actin filament velocity in root epidermal cells confirmed its reduction in *rhd2-1* GFP-FABD2 transgenic line as compared to Col-0 GFP-FABD2 (Figure 8E; Supplementary Movie S1).

**Figure 8.**
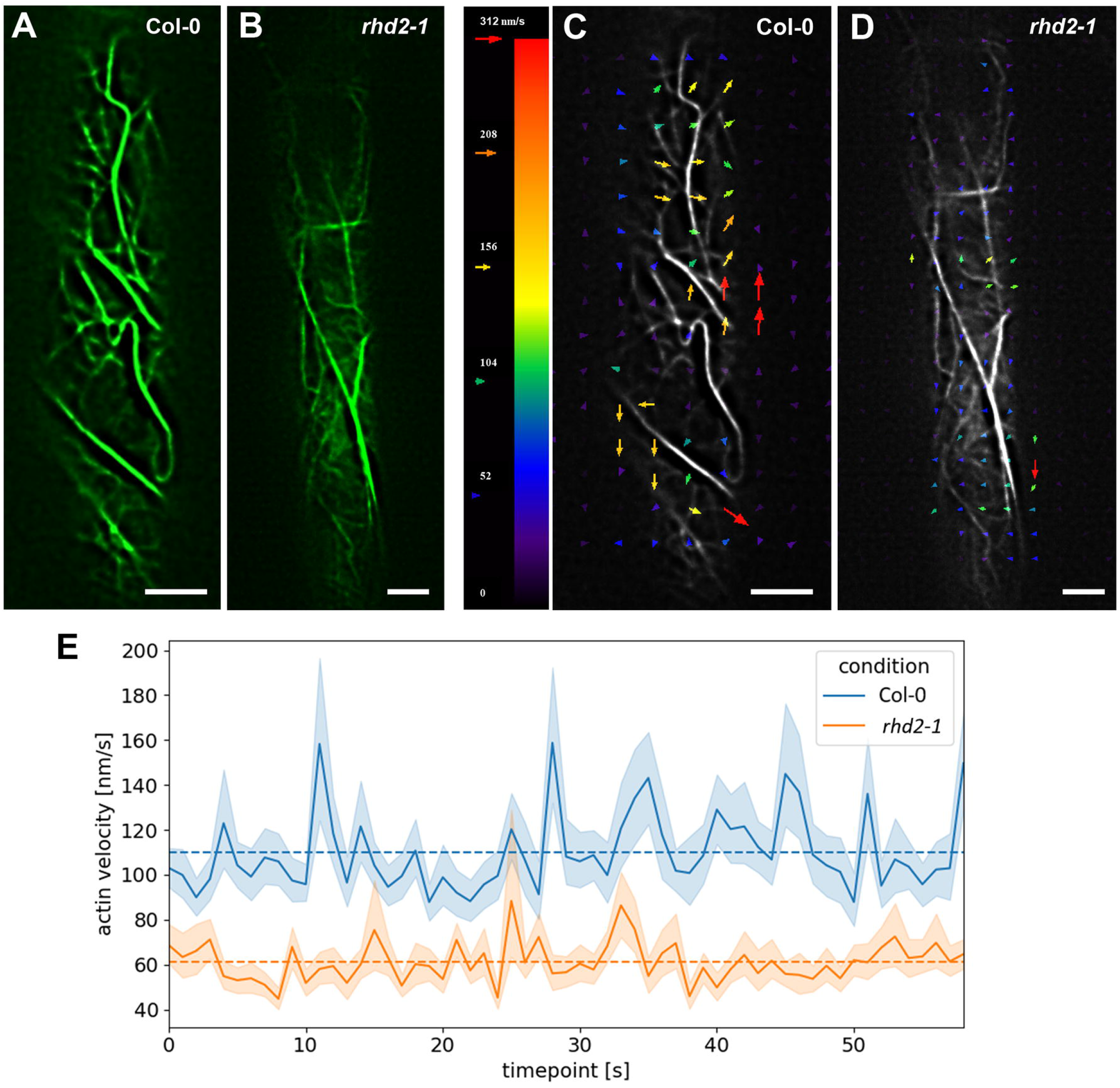
Analysis of actin velocity in root cells of Col-0 and *rhd2-1* using lattice light-sheet fluorescence microscopy. (A, B) Actin cytoskeleton in epidermal root cells of transgenic lines Col-0 with GFP-FABD2 (A) and *rhd2-1* with GFP-FABD2 (B). (C, D) Velocity of actin filaments in root epidermal cells from PIV analyser plugin (Fiji) of the Col-0 with GFP-FABD2 (C) and *rhd2-1* with GFP-FABD2 transgenic lines (D). Mean actin velocity from 60 s long recording in root epidermal cells of the Col-0 with GFP-FABD2 and *rhd2-1* with GFP-FABD2 transgenic lines (E). Measurements are from 7 cells of 5 plants in Col-0, and 5 cells of 2 plants in *rhd2-1* mutant. Scale bars = 5 µm.

In summary, loss-of-function mutation in *AtRBOHC/RHD2* gene caused pleiotropic proteome-wide effects on plant development and altered drought stress response. It was likely caused by altered defective root-to-shoot signalling, membrane and vesicular transport, microtubule organisation and actin filament dynamics.

## Discussion

Differential proteomic analysis revealed that functional mutation in *RBOHC/RHD2* affected PM-localised aquaporins (PIPs). These proteins facilitate water membrane diffusion and regulate the root hydraulic conductivity (Maurel *et al*., 2015). PIPs are implicated in plant abiotic and biotic stress responses (Afzal *et al*., 2016) and their genetic modification substantially modulates the plant stress response (Zhou *et al*., 2014). They are subjected to constant endocytic recycling that is experimentally proved mostly on PIP2 isoform (Besserer *et al*., 2012; Martinière *et al*., 2019). After biosynthesis in the endoplasmic reticulum, they are transported via Golgi apparatus to the *trans*-Golgi network (TGN), and might either be delivered to PM, recycled, or directed to the multivesicular body (MVB) for degradation (Hachez *et al*., 2013; Chevalier and Chaumont, 2015). The dynamic cycling of AtPIP2;1 in control conditions through the constitutive endocytosis from the PM to internal compartments is executed by clathrin-mediated endocytosis (Li *et al*., 2011). The cycle starts by endocytosis, proceeds through transfer to the TGN, and further recycling back to the PM (Chevalier and Chaumont, 2015). The insertion of PIPs into PM occurs via a SNARE complex (Besserer *et al*., 2012; Hachez *et al*., 2014), and is regulated by homo- and heteromerisation (Otto *et al*., 2010) as well as posttranslational regulation of PIPs (Santoni, 2017). PIPs are internalised to cells via clathrin-mediated or clathrin-independent lipid raft-associated endocytosis (Hachez *et al*., 2014). Moreover, subcellular trafficking of PIPs is sensitive to external conditions and its disruption alters the membrane water permeability (Wudick *et al*., 2015).

Arabidopsis *rhd2-1* mutant dramatically differed from Col-0 in proteins involved in vesicular transport including ESCRT, SNARE, COG, exocyst and retromer complexes. ESCRT complex is involved in MVB biogenesis, but ESCRT proteins are also essential for protein recycling from endosomes to the PM, sorting ubiquitinated cargoes into lysosomes/vacuoles, exosome secretion and autophagy (Gao *et al*., 2017). ESCRT-I component ELC, also named as VPS23 (2.9 fold upregulated in the *rhd2-1* mutant) affects the subcellular localisation of SALT OVERLY SENSITIVE 2 (SOS2) to the PM, thus essentially modulating plant salt stress tolerance (Lou *et al*., 2020).

Retromer recycles vacuolar sorting receptors from the TGN (Niemes *et al*., 2010). It also mediates the retrograde recycling of biosynthetic cargo receptors back to the TGN/Golgi and endocytic cargos back to the PM (González Solís *et al*., 2022). Retromer subunits VPS29 (1.5 fold upregulated in the *rhd2-1* mutant) and VPS35A are important for prevacuolar compartment (PVC) function in Arabidopsis (Nodzyński *et al*., 2013). VPS29 is also required for endosome homeostasis, PIN protein cycling, and dynamic PIN1 repolarisation during development (Jaillais *et al*., 2007).

SNAREs mediate membrane fusions and confer membrane fusion specificity. Qa-SNARE SYP22 (downregulated in the *rhd2-1* mutant) is involved in the vacuolar transport pathway of Arabidopsis and is localised to prevacuolar compartments and tonoplast (Uemura *et al*., 2004). SYP22, together with VAMP727, regulate BRI1 targeting to PM, contributing to plant growth in Arabidopsis (Zhang *et al*., 2019). Another protein differentially regulated in the *rhd2-1* mutant, involved in PM targeting, was PROPHOSPHOINOSITIDE PHOSPHATASE SAC8 (downregulated in *rhd2-1* roots). It is required for PtdIns4P, PtdIns(4,5)P2 homeostasis and is implicated in trafficking of the PM-localised proteins PIN1 and PIN2 from TGN back to PM (Song *et al*., 2021). The role of PtdIns4P and PtdIns(4,5)P2 in the internalisation of AtPIP2;1 from the PM to the vacuole under salt stress was linked to phosphatidylinositol 3-kinase and phosphatidylinositol 4-kinase. This internalisation proceeded through clathrin-mediated endocytosis (Ueda *et al*., 2016). The process of AtPIP2;1 recycling from the PM via clathrin-mediated endocytosis under control conditions may, however, switch to clathrin-independent endocytosis induced by salt stress (Luu *et al*., 2012). Such significant alterations and defects in vesicular transport might be a reason of a substantially changed abundance of PM and tonoplast-localised proteins in the *rhd2-1* mutant, including the PIP1 isoforms. It is likely that loss of AtRBOHC/RHD2 function might essentially affect the protein delivery to the PM and tonoplast, leading to diverse physiological consequences.

The mechanism linking ROS-producing activities of RBOHC to such effects is elusive. Endocytosis of PIP2-1 depends on ROS produced by RBOHD and its clustering during salt stress in Arabidopsis (Martinière *et al*., 2019). Although a similar regulatory mechanism might be assigned also to RBOHC, it hardly can be generalised to all proteins found in this proteomic screen. In agreement, PM-localised proton ATPase (PM H^+^-ATPase) AHA2 showed a RBOHD-independent pattern of recycling during osmotic stress (Martinière *et al*., 2019).

Our results indicate that apart from root hair growth regulation, the production of ROS by AtRBOHC/RHD2 is likely important for the amount and stability of PIP1 isoforms, affecting the Arabidopsis drought stress response. It has been shown that endocytic internalisation of PIPs into intracellular compartments under abiotic stresses such as drought, was directly related to the effort of plants to downregulate root water transport due to water deprivation under stress conditions. The rate of exchange between the PM and intracellular compartments of AtPIP1;2 and AtPIP2;1 revealed that the aquaporin recycling was greater under salt stress than under control conditions (Luu *et al*., 2012). This may reflect also differences at the shoot-to-root ratio of PIPs abundance level. Studies in Arabidopsis revealed discrepancies between the level of mRNA expression and the protein abundances of PIPs. Frequently after drought stress, a higher PIPs protein levels were observed in shoots and less in roots (Yepes-Molina *et al*., 2020). In addition, it has been shown that under drought stress, the expression level of AtPIP2;5 was increased in the aerial parts of Arabidopsis plants (Jang *et al*., 2004). Nevertheless, the drought stress sensitivity of *rhd2-1* mutant is most likely determined also by other PM residing proteins found in the proteomic screen, such as PM H^+^-ATPase AHA2 (Pei *et al*., 2022), CASPARIAN STRIP MEMBRANE DOMAIN PROTEIN 1 (CASP1) (Roppolo *et al*., 2014; Kamiya *et al*., 2015), and ABCG32 (Seo and Park, 2011).

In conclusion, proteomic analysis uncovered, that the functional mutation in *RBOHC/RHD2* leads to substantial defects in vesicular transport, affecting homeostasis of proteins residing at the PM, and alteration of plant drought stress response.

Microtubules are fundamental components of the cytoskeleton inevitable for cell division, growth and morphogenesis. They also perceive signals derived from external environment, including mechanical stimuli, ROS, abiotic and biotic stresses (Nick *et al*., 2013). Microtubules in stomata guard cells regulate stomatal closure and water loss by controlling AtPIP2;1 dynamic behaviour at PM (Cui et al., 2021). Our results point to the dependence of PIP1 aquaporin isoforms abundances on root specific *AtRBOHC/RHD2* expression, and the concomitant increased levels of cortical microtubule isotropy in *rhd2-1* mutant imply that microtubules may mediate the AtRBOHC/RHD2-dependent PIP1 dynamics. It has been shown that in the roots of the halophyte ice plant (*Mesembryanthemum crystallinum* L.) under salinity stress conditions abundance of PIPs at the PM was altered. It was demonstrated that the amount of McPIP1;4 and McPIP2;1 increased at the PM under salinity conditions and their trafficking was performed by clathrin-coated vesicles (CCV). The proteomic analysis from positively charged microsome fractions revealed McPIPs interactions with CCV-associated proteins, such as the CLATHRIN HEAVY CHAIN (CHC), further confirmed by CCV enrichment assays and immunoblotting using anti-CHC antibody (Gómez-Méndez *et al*., 2023). Most likely, the microtubule reorientation in *rhd2-1* mutant is related to the altered ROS production, as chemical modulation of ROS was reported to alter their integrity (Livanos *et al*., 2014). These alterations in cortical microtubule dynamic properties and orientation are likely associated with an increased abundance of yet uncharacterised MICROTUBULE ASSOCIATED PROTEIN 70-2 (MAP70-2), belonging to a plant-specific MAP70 family (Korolev *et al*., 2005). Further, increased abundances of two TUBULIN isoforms may indicate their altered turnover in the *rhd2-1* mutant.

Our results show that AtRBOHC/RHD2 may be important for actin dynamics as well. The link between RBOH and actin was established previously showing that ROS generated by RBOHD is important for actin rearrangements during innate immunity (Li *et al*., 2017; Cao *et al*., 2022). Additional work revealed that FLS2 endocytosis was inhibited by the myosin inhibitor 2,3-butanedione monoxime, as well as treatment with Lat B. These results suggest that the ligand-mediated endocytosis of pathogenesis-related receptors requires the actin cytoskeleton (Beck *et al*., 2012). However, in comparison to other systems like yeast, actin participation has not been proved at the spots of clathrin-mediated endocytosis in plants. Actin is, however, involved in further delivery of post-endocytic vesicles, for their intracellular movement, interaction with endosomal compartments, and for cargo sorting (Aniento *et al*., 2022; Narasimhan *et al*., 2020). Actin cytoskeleton and actin-associated proteins were already linked to plant stress reactions. Stomatal closure during drought stress is regulated by ABA signalling that regulates actin reorganization (Shi *et al*., 2022). It has been shown that SWR1 (SWI2/SNF2-RELATED CHROMATIN REMODELLING 1) complex is involved in positive regulation of the expression levels of stress-responsive *ABA INSENSITIVE 1* (*ABI1*) and *ABI2* genes. Interestingly, ACTIN-RELATED PROTEIN 6 (ARP6) is one component of this complex and in line with that, the *arp6* mutant was more sensitive to salt stress. Transcript levels of *ABI1* and *ABI2* genes were significantly lower in *arp6* mutant after salt treatment (Do *et al*., 2023). Cytosolic ABA Receptor Kinase 3 (CARK3) is involved in phosphorylation of ABA receptors, mediating ABA signalling pathway. CARK3 interacts with ACTIN DEPOLYMERIZATION FACTOR 4 (ADF4) and regulates it by phosphorylation. Interestingly, overexpression of ADF4 enhanced drought resistance of tested plants and soil drought tolerance assay revealed that the survival of *adf4* mutant was significantly lower in comparison to control plants after drought treatment (Peng *et al*., 2022). The role of ADF4 was approved also in stomatal closure, which was mediated through reorganisation of actin filaments (Zhao *et al*., 2016). Other study revealed ADF5-related F-actin-bundling activity, which was positively involved in regulation of drought stress response. This was documented by increased drought stress sensitivity of *adf5* mutant (Qian *et al*., 2019). Similarly, cotton GhVLN4 responsible for actin microfilament bundling, upon overexpression in Arabidopsis, conferred resistance to *Verticilium dahliae*, salt and drought stresses (Ge *et al*., 2021). As demonstrated by our results, AtRBOHC/RHD2 may be another RBOH isoform mediating actin dynamics in Arabidopsis, possibly influencing also the phenotypical changes in root hair development of *rhd2-1* mutant. It is likely, that the altered ROS production might contribute to modification of actin dynamics involved in the root hair phenotype of *rhd2-1* mutant. Lower actin dynamics is most likely linked to the altered endosomal motility in the *rhd2-1* mutant. Indeed, movements of TGN/early endosomes visualised with GFP-RabA1d were affected in the bulging domains of the *rhd2-1* mutant (Kuběnová *et al*., 2022). Here we show that early endosomal protein BREFELDIN A-INHIBITED GUANINE NUCLEOTIDE-EXCHANGE PROTEIN 5 (BIG5; Tanaka *et al*., 2009), usually abundant in the primary root differentiation zone (Table 1), was downregulated in the *rhd2-1* mutant and might contribute to its root hair phenotype.

All in all, altered abundances of PIPs, CLATHRIN, ESCRT, SNAREs along with cortical microtubule reorganisation might significantly contribute to the developmental phenotype of *rhd2-1* and its higher sensitivity to drought stress.

## Author contributions

TT, LK, OŠ, PD, JH and STB performed the experiments and analyses. JŠ coordinated the experiments, provided funding, supervised the project and helped with data assessment. JŠ and PT provided the infrastructure and helped with the interpretation of the results. TT, LK, OŠ, PD and MO drafted the manuscript which was improved and edited by TT, MO and JŠ. All authors approved the final version of the manuscript.

## Funding statement

This work was supported by the Czech Science Foundation GACR (project Nr. 19-18675S).

## Acknowledgement

Mass spectrometry analyses were performed at the Turku Proteomics Facility supported by Biocentre Finland.

## Conflict of interest

The authors declare that they have no conflict of interest.

## Data Availability

The mass spectrometry proteomics data have been deposited to the ProteomeXchange Consortium via the PRIDE (Perez-Riverol *et al*., 2022) partner repository with the dataset identifier PXD038577”.

## Notes

### Competing Interest Statement

The authors have declared no competing interest.

## References

Afzal Z, Howton T, Sun Y, Mukhtar M. 2016. The roles of aquaporins in plant stress responses. Journal of Developmental Biology 4, 9.

Aniento F, Sánchez de Medina Hernández V, Dagdas Y, Rojas-Pierce M, Russinova E. 2022. Molecular mechanisms of endomembrane trafficking in plants. Plant Cell 34, 146–173.

Arenas-Alfonseca L, Gotor C, Romero LC, García I. 2018. Role of mitochondrial cyanide detoxification in Arabidopsis root hair development. Plant Signaling & Behavior 13, e1537699.

Beck M, Zhou J, Faulkner C, MacLean D, Robatzek S. 2012. Spatio-temporal cellular dynamics of the Arabidopsis flagellin receptor reveal activation status-dependent endosomal sorting. Plant Cell 24, 4205–4219.

Besserer A, Burnotte E, Bienert GP, Chevalier AS, Errachid A, Grefen C, Blatt MR, Chaumont F. 2012. Selective regulation of maize plasma membrane aquaporin trafficking and activity by the SNARE SYP121. Plant Cell 24, 3463–3481.

Cao L, Wang W, Zhang W, Staiger CJ. 2022. Lipid signaling requires ROS production to elicit actin cytoskeleton remodeling during plant innate immunity. International Journal of Molecular Sciences 23, 2447.

Carol R, Takeda S, Linstead P, Durrant MC, Kakesova H, Derbyshire P, Drea S, Zarsky V, Dolan L. 2005. A RhoGDP dissociation inhibitor spatially regulates growth in root hair cells. Nature 438, 1013–1016.

Chapman JM, Muhlemann JK, Gayomba SR, Muday GK. 2019. RBOH-dependent ROS synthesis and ROS scavenging by plant specialized metabolites to modulate plant development and stress responses. Chemical Research in Toxicology 32, 370–396.

Chen BC, Legant WR, Wang K et al. 2014. Lattice light-sheet microscopy: imaging molecules to embryos at high spatiotemporal resolution. Science 346, (6208):1257998.

Chevalier AS, Chaumont F. 2015. Trafficking of plant plasma membrane aquaporins: multiple regulation levels and complex sorting signals. Plant & Cell Physiology 56, 819–829.

Chronopoulou E, Georgakis N, Nianiou-Obeidat I, Madesis P, Perperopoulou F, Pouliou F, Vasilopoulou E, Ioannou E, Ataya FS, Labrou NE. 2017. Plant glutathione transferases in abiotic stress response and herbicide resistance. In: Hossain MA, Mostofa MG, Diaz-Vivancos P, Burritt DJ, Fujita M, Tran L-SP, eds. Glutathione in plant growth, development, and stress tolerance. Springer International Publishing, 215–233.

Cui Y, Zhao Y, Lu Y, Su X, Chen Y, Shen Y, Lin J, Li X. 2021. In vivo single-particle tracking of the aquaporin AtPIP2;1 in stomata reveals cell type-specific dynamics. Plant Physiology 185, 1666–1681.

Do BH, Hiep NT, Lao TD, Nguyen NH. 2023. Loss-of-function mutation of ACTIN-RELATED PROTEIN 6 (ARP6) impairs root growth in response to salinity stress. Molecular Biotechnology, https://doi.org/10.1007/s12033-023-00653-x

Foreman J, Demidchik V, Bothwell JH, Mylona P, Miedema H, Torres MA, Linstead P, Costa S, Brownlee C, Jones JD, Davies JM, Dolan L. 2003. Reactive oxygen species produced by NADPH oxidase regulate plant cell growth. Nature 422, 442–446.

Fucile G, Di Biase D, Nahal H, La G, Khodabandeh S, Chen Y, Easley K, Christendat D, Kelley L, Provart NJ. 2011. ePlant and the 3D data display initiative: integrative systems biology on the world wide web. PLoS One 6(1):e15237.

Fujita M, Himmelspach R, Hocart CH, Williamson RE, Mansfield SD, Wasteneys GO. 2021. Cortical microtubules optimise cell-wall crystallinity to drive unidirectional growth in Arabidopsis. Plant Journal 66, 915–928.

Gao C, Zhuang X, Shen J, Jiang L. 2017. Plant ESCRT Complexes: Moving Beyond Endosomal Sorting. Trends in Plant Science 22, 986–998.

Ge D, Pan T, Zhang P, Wang L, Zhang J, Zhang Z, Dong H, Sun J, Liu K, Lv F. 2021. GhVLN4 is involved in multiple stress responses and required for resistance to verticillium wilt. Plant Science 302, 110629.

Gómez-Méndez MF, Amezcua-Romero JC, Rosas-Santiago P, Hernández-Domínguez EE, de Luna-Valdez LA, Ruiz-Salas JL, Vera-Estrella R, Pantoja O. 2023. Ice plant root plasma membrane aquaporins are regulated by clathrin-coated vesicles in response to salt stress. Plant Physiology 191, 199–218.

González Solís A, Berryman E, Otegui MS. 2022. Plant endosomes as protein sorting hubs. FEBS Letters 596, 2288–2304.

Hachez C, Besserer A, Chevalier AS, Chaumont F. 2013. Insights into plant plasma membrane aquaporin trafficking. Trends in Plant Science 18, 344–352.

Hachez C, Laloux T, Reinhardt H, Cavez D, Degand H, Grefen C, De Rycke R, Inzé D, Blatt MR, Russinova E, Chaumont F. 2014. *Arabidopsis* SNAREs SYP61 and SYP121 coordinate the trafficking of plasma membrane aquaporin PIP2;7 to modulate the cell membrane water permeability. Plant Cell 26, 3132–3147.

Hao H, Fan L, Chen T, Li R, Li X, He Q, Botella MA, Lin J. 2014. Clathrin and membrane microdomains cooperatively regulate RBOH D dynamics and activity in Arabidopsis. Plant Cell 26, 1729–1745.

Jaillais Y, Santambrogio M, Rozier F, Fobis-Loisy I, Miège C, Gaude T. 2007. The retromer protein VPS29 links cell polarity and organ initiation in plants. Cell 130, 1057–1070.

Jang JY, Kim DG, Kim YO, Kim JS, Kang H. 2004. An expression analysis of a gene family encoding plasma membrane aquaporins in response to abiotic stresses in Arabidopsis thaliana. Plant Molecular Biology 54, 713–725.

Jones MA, Raymond MJ, Yang Z, Smirnoff N. 2007. NADPH oxidase-dependent reactive oxygen species formation required for root hair growth depends on ROP GTPase. Journal of Experimental Botany 58, 1261–1270.

Jones MA, Raymond MJ, Smirnoff N. 2006. Analysis of the root-hair morphogenesis transcriptome reveals the molecular identity of six genes with roles in root-hair development in Arabidopsis. Plant Journal 45(1), 83–100.

Kamiya T, Borghi M, Wang P, Danku JMC, Kalmbach L, Hosmani PS, Naseer S, Fujiwara T, Geldner N, Salt DE. 2015. The MYB36 transcription factor orchestrates Casparian strip formation. Proceedings of the National Academy of Sciences 112, 10533– 10538.

Kaur G, Pati PK. 2018. In silico insights on diverse interacting partners and phosphorylation sites of respiratory burst oxidase homolog (Rbohs) gene families from Arabidopsis and rice. BMC Plant Biology 18, 161.

Kaya H, Takeda S, Kobayashi MJ, Kimura S, Iizuka A, Imai A, Hishinuma H, Kawarazaki T, Mori K, Yamamoto Y, Murakami Y, Nakauchi A, Abe M, Kuchitsu K. 2019. Comparative analysis of the reactive oxygen speciesCproducing enzymatic activity of Arabidopsis NADPH oxidases. Plant Journal 98, 291–300.

Korolev AV, Chan J, Naldrett MJ, Doonan JH, Lloyd CW. 2005. Identification of a novel family of 70 kDa microtubule-associated proteins in Arabidopsis cells: Novel family of plant MAPs. Plant Journal 42, 547–555.

Kuběnová L, Tichá M, Šamaj J, Ovečka M. 2022. ROOT HAIR DEFECTIVE 2 vesicular delivery to the apical plasma membrane domain during Arabidopsis root hair development. Plant Physiology 188, 1563–1585.

Li J, Cao L, Staiger CJ. 2017. Capping protein modulates actin remodeling in response to reactive oxygen species during plant innate immunity. Plant Physiology 173, 1125–1136.

Li X, Wang X, Yang Y, Li R, He Q, Fang X, Luu DT, Maurel C, Lin J. 2011. Single-molecule analysis of PIP2;1 dynamics and partitioning reveals multiple modes of Arabidopsis plasma membrane aquaporin regulation. Plant Cell 23, 3780–3797.

Livanos P, Galatis B, Apostolakos P. 2014. The interplay between ROS and tubulin cytoskeleton in plants. Plant Signaling & Behavior 9: e28069.

Lou L, Yu F, Tian M, Liu G, Wu Y, Wu Y, Xia R, Pardo JM, Guo Y, Xie Q. 2020. ESCRT-I component VPS23A sustains salt tolerance by strengthening the SOS module in Arabidopsis. Molecular Plant 13, 1134–1148.

Luu DT, Martiniãre A, Sorieul M, Runions J, Maurel C. 2012. Fluorescence recovery after photobleaching reveals high cycling dynamics of plasma membrane aquaporins in Arabidopsis roots under salt stress. Plant Journal 69, 894–905.

Mangano S, Denita-Juarez SP, Marzol E, Borassi C, Estevez JM. 2018. High auxin and high phosphate impact on RSL2 expression and ROS-homeostasis linked to root hair growth in *Arabidopsis thaliana*. Frontiers in Plant Science 9, 1164.

Martin RE, Marzol E, Estevez JM, Muday GK. 2022. Ethylene signaling increases reactive oxygen species accumulation to drive root hair initiation in *Arabidopsis*. Development 149, dev200487.

Martinière A, Fiche JB, Smokvarska M, Mari S, Alcon C, Dumont X, Hematy K, Jaillais Y, Nollmann M, Maurel C. 2019. Osmotic stress activates two reactive oxygen species pathways with distinct effects on protein nanodomains and diffusion. Plant Physiology 179, 1581–1593.

Maurel C, Boursiac Y, Luu D-T, Santoni V, Shahzad Z, Verdoucq L. 2015. Aquaporins in plants. Physiological Reviews 95, 1321–1358.

Melicher P, Dvořák P, Krasylenko Y, Shapiguzov A, Kangasjärvi J, Šamaj J, Takáč T. 2022. Arabidopsis iron superoxide dismutase FSD1 protects against methyl viologen-induced oxidative stress in a copper-dependent manner. Frontiers in Plant Science 13, 823561.

Monshausen GB, Bibikova TN, Messerli MA, Shi C, Gilroy S. 2007. Oscillations in extracellular pH and reactive oxygen species modulate tip growth of *Arabidopsis* root hairs. Proceedings of the National Academy of Sciences 104, 20996–21001.

Narasimhan M, Johnson A, Prizak R, Kaufmann WA, Tan S, Casillas-Pérez B, Friml J. 2020. Evolutionarily unique mechanistic framework of clathrin-mediated endocytosis in plants. eLife 9, e52067.

Nick P. 2013. Microtubules, signalling and abiotic stress. Plant Journal 75, 309–323.

Niemes S, Langhans M, Viotti C, Scheuring D, San Wan Yan M, Jiang L, Hillmer S, Robinson DG, Pimpl P. 2010. Retromer recycles vacuolar sorting receptors from the trans-Golgi network. Plant Journal 61, 107–121.

Nodzyński T, Feraru MI, Hirsch S, De Rycke R, Niculaes C, Boerjan W, Van Leene J, De Jaeger G, Vanneste S, Friml J. 2013. Retromer Subunits VPS35A and VPS29 Mediate Prevacuolar Compartment (PVC) Function in Arabidopsis. Molecular Plant 6, 1849–1862.

Otto B, Uehlein N, Sdorra S et al. 2010. Aquaporin Tetramer Composition Modifies the Function of Tobacco Aquaporins. Journal of Biological Chemistry 285, 31253–31260.

Pasternak T, Tietz O, Rapp K, Begheldo M, Nitschke R, Ruperti B, Palme K. 2015. Protocol: an improved and universal procedure for whole-mount immunolocalisation in plants. Plant Methods 11, 50.

Pei D, Hua D, Deng J, Wang Z, Song C, Wang Y, Wang Y, Qi J, Kollist H, Yang S, Guo Y, Gong Z. 2022. Phosphorylation of the plasma membrane H+-ATPase AHA2 by BAK1 is required for ABA-induced stomatal closure in Arabidopsis. Plant Cell 34, 2708–2729.

Peng L, He J, Yao H, Yu Q, Zhang Q, Li K, Huang Y, Chen L, Li X, Yang Y, Li X. 2022. CARK3-mediated ADF4 regulates hypocotyl elongation and soil drought stress in Arabidopsis. Frontiers in Plant Science 13, 1065677.

Perez-Riverol Y, Bai J, Bandla C, Hewapathirana S, García-Seisdedos D, Kamatchinathan S, Kundu D, Prakash A, Frericks-Zipper A, Eisenacher M, Walzer M, Wang S, Brazma A, Vizcaíno JA 2022. The PRIDE database resources in 2022: A Hub for mass spectrometry-based proteomics evidences. Nucleic Acids Res 50(D1):D543–D552.

Qian D, Zhang Z, He J, Zhang P, Ou X, Li T, Niu L, Nan Q, Niu Y, He W, An L, Jiang K, Xiang Y. 2019. Arabidopsis ADF5 promotes stomatal closure by regulating actin cytoskeleton remodeling in response to ABA and drought stress. Journal of Experimental Botany 70, 435–446.

Roppolo D, Boeckmann B, Pfister A, Boutet E, Rubio MC, Dénervaud-Tendon V, Vermeer JEM, Gheyselinck J, Xenarios I, Geldner N. 2014. Functional and evolutionary analysis of the CASPARIAN STRIP MEMBRANE DOMAIN PROTEIN family. Plant Physiology 165, 1709–1722.

Santoni V. 2017. Plant Aquaporin Posttranslational Regulation. In: Chaumont F, Tyerman SD, eds. Plant Aquaporins: from transport to signaling. Cham: Springer International Publishing, 83–105.

Šamajová O, Komis G, Šamaj J. 2014. Immunofluorescent localisation of MAPKs and colocalisation with microtubules in Arabidopsis seedling whole-mount probes. Methods in Molecular Biology 1171, 107–115.

Shi Y, Liu X, Zhao S, Guo Y. 2022. The PYR-PP2C-CKL2 module regulates ABA-mediated actin reorganization during stomatal closure. New Phytologist 233, 2168–2184.

Schiefelbein JW, Somerville C. 1990. Genetic control of root hair development in *Arabidopsis thaliana*. Plant Cell 2, 235–243.

Schindelin J, Arganda-Carreras I, Frise E et al. 2012 Fiji: an open-source platform for biological-image analysis. Nature Methods 9, 676 – 682.

Szklarczyk D, Gable AL, Lyon D et al. 2019. STRING v11: protein-protein association networks with increased coverage, supporting functional discovery in genome-wide experimental datasets. Nucleic Acids Research 47(D1), D607–D613.

Seo PJ, Park C-M. 2011. Cuticular wax biosynthesis as a way of inducing drought resistance. Plant Signaling & Behavior 6, 1043–1045.

Smokvarska M, Francis C, Platre MP et al. 2020. A plasma membrane nanodomain ensures signal specificity during osmotic signaling in plants. Current Biology 30, 4654–4664.

Song L, Wang Y, Guo Z, Lam SM, Shui G, Cheng Y. 2021. NCP2/RHD4/SAC7, SAC6 and SAC8 phosphoinositide phosphatases are required for PtdIns4P and PtdIns(4,5)P2 homeostasis and Arabidopsis development. New Phytologist 231, 713–725.

Takáč T, Šamajová O, Pechan T, Luptovčiak I, Šamaj J. 2017. Feedback microtubule control and microtubule-actin cross-talk in Arabidopsis Revealed by integrative proteomic and cell biology analysis of KATANIN 1 mutants. Molecular & Cellular Proteomics 16, 1591–1609.

Tanaka H, Kitakura S, De Rycke R, De Groodt R, Friml J. 2009. Fluorescence imaging-based screen identifies ARF GEF component of early endosomal trafficking. Current Biology 19, 391–397.

Ueda M, Tsutsumi N, Fujimoto M. 2016. Salt stress induces internalization of plasma membrane aquaporin into the vacuole in *Arabidopsis thaliana*. Biochemical and Biophysical Research Communications 474, 742–746.

Uemura T, Ueda T, Ohniwa RL, Nakano A, Takeyasu K, Sato MH. 2004. Systematic analysis of SNARE molecules in Arabidopsis: dissection of the post-Golgi network in plant cells. Cell Structure and Function 29, 49–65.

Wudick MM, Li X, Valentini V, Geldner N, Chory J, Lin J, Maurel C, Luu D-T. 2015. Subcellular redistribution of root aquaporins induced by hydrogen peroxide. Molecular Plant 8, 1103–1114.

Yepes-Molina L, Bárzana G, Carvajal M. 2020. Controversial regulation of gene expression and protein transduction of aquaporins under drought and salinity stress. Plants 9, 1662.

Zhao S, Jiang Y, Zhao Y, Huang S, Yuan M, Zhao Y, Guo Y. 2016. Casein Kinase1-like Protein 2 regulates actin filament stability and stomatal closure via phosphorylation of actin depolymerizing factor. Plant Cell 28, 1422–1439.

Zhang X, Köster P, Schlücking K, Balcerowicz D, Hashimoto K, Kuchitsu K, Vissenberg K, Kudla J. 2018. CBL1CCIPK26Cmediated phosphorylation enhances activity of the NADPH oxidase RBOHC, but is dispensable for root hair growth. FEBS Letters 592, 2582– 2593.

Zhang L, Liu Y, Zhu XF, Jung JH, Sun Q, Li TY, Chen LJ, Duan YX, Xuan YH. 2019. SYP22 and VAMP727 regulate BRI1 plasma membrane targeting to control plant growth in Arabidopsis. New Phytologist 223, 1059–1065.

Zhou L, Wang C, Liu R, Han Q, Vandeleur RK, Du J, Tyerman S, Shou H. 2014. Constitutive overexpression of soybean plasma membrane intrinsic protein GmPIP1;6 confers salt tolerance. BMC Plant Biology 14, 181.

